# SPEAQeasy: a Scalable Pipeline for Expression Analysis and Quantification for R/Bioconductor-powered RNA-seq analyses

**DOI:** 10.1101/2020.12.11.386789

**Authors:** Nicholas J. Eagles, Emily E. Burke, Jacob Leonard, Brianna K. Barry, Joshua M. Stolz, Louise Huuki, BaDoi N. Phan, Violeta Larios Serrato, Everardo Gutiérrez-Millán, Israel Aguilar-Ordoñez, Andrew E. Jaffe, Leonardo Collado-Torres

## Abstract

RNA sequencing (RNA-seq) is a common and widespread biological assay, and an increasing amount of data is generated with it. In practice, there are a large number of individual steps a researcher must perform before raw RNA-seq reads yield directly valuable information, such as differential gene expression data. Existing software tools are typically specialized, only performing one step-- such as alignment of reads to a reference genome-- of a larger workflow. The demand for a more comprehensive and reproducible workflow has led to the production of a number of publicly available RNA-seq pipelines. However, we have found that most require computational expertise to set up or share among several users, are not actively maintained, or lack features we have found to be important in our own analyses. In response to these concerns, we have developed a Scalable Pipeline for Expression Analysis and Quantification (SPEAQeasy), which is easy to install and share, and provides a bridge towards R/Bioconductor downstream analysis solutions. SPEAQeasy is user-friendly and lowers the computational-domain entry barrier for biologists and clinicians to RNA-seq data processing as the main input file is a table with sample names and their corresponding FASTQ files. SPEAQeasy is portable across computational frameworks (SGE, SLURM, local, docker integration) and different configuration files are provided.

## Introduction

Gene expression analyses have been revolutionized by the emergence of high-throughput sequencing (*1–3*) which has enabled an explosion in RNA-sequencing (RNA-seq) projects (*4–6*). Sequencing machines typically output the data in the FASTQ format (*7*) that can amount to several gigabytes of disk space per sample depending on the read length and coverage depth of a given experiment. Before doing any statistical analyses on this data such as differential expression (*8, 9*), researchers need to process the GBs or even terabytes of data to compress it and extract the desired information. Doing so requires computationally demanding steps such as RNA-seq alignment (*10–12*) and read quantification (*13, 14*). Since the emergence of RNA-seq, a diverse set of bioinformatics software has been designed to solve specific steps of the RNA-seq processing (*15–17*).

Several RNA-seq processing bioinformatics pipelines have been developed to tie these required processing steps together (*18–23*). The common goal of these approaches involves helping biologists and researchers weave together these bioinformatics solutions to uniformly process samples from RNA-seq projects with different characteristics; for example, single-end versus paired-end. RNA-seq processing pipelines have different characteristics such as the RNA-seq aligner of choice and the quality control steps they use. The design choices of each RNA-seq processing pipeline can have an impact on which analyses researchers can perform with the processed data. Furthermore, the ease of software installation, portability, and level of support can affect the usability of these pipelines.

In recent years we have worked on several RNA-seq projects (*24–26*) and designed an RNA-seq processing pipeline that satisfied our needs to generate quality checked and uniformly processed data with several quality control metrics we could then use in our statistical analyses. We then improved the usability and portability of this pipeline thanks to the Nextflow framework (*27*). Our solution, SPEAQeasy, ultimately generates `RangedSummarizedExperiment` R objects (*28*) that are the foundation block for many Bioconductor R packages and the statistical methods they provide (*8, 9, 29, 30*). Other key features of SPEAQeasy are that it produces the information that coupled with DNA genotyping information can be used for detecting and fixing sample swaps (*31–33*), RNA-seq processing quality metrics that are helpful for statistically adjusting for quality differences across samples (*5*), data that powers the exploration of the un-annotated transcriptome, and that it can be used in several computational frameworks thanks to Nextflow’s configuration flexibility (*27*).

## Results

### Overview

We have developed a portable RNA sequencing (RNA-seq) processing pipeline, SPEAQeasy, that provides analysis-ready gene expression files (**Figure 2**). SPEAQeasy is a Nextflow-powered (*27*) pipeline that starts from a set of FASTQ files (*7*), performs quality assessment and other processing steps (Methods: overview), and produces easy-to-use R objects (*29*). SPEAQeasy facilitates both traditional RNA-seq downstream analyses such as gene differential expression, but also the exploration of the annotated transcriptome (*34, 35*) by quantifying reads that span exon-exon junctions and providing bigWig base-pair coverage files (*36*). Input RNA-seq reads are aligned using HISAT2 (*37*) to a reference genome and pseudo-aligned to a reference transcriptome using kallisto (*38*) or Salmon (*39*). Genes, exons, and exon-exon junctions are then quantified using featureCounts (*14*) and regtools (*40*). The resulting quality metrics and read quantification outputs are then arranged to create `summarizedExperiment` (*28*) R objects that combine the read quantification, expression feature information, and processing and quality metrics. These SummarizedExperiment objects can then be used with a wide variety of Bioconductor (*29*) R packages to perform downstream analyses such as differential expression (*8, 9, 30*), identification of differentially expressed regions (DER) (*41*), and exploratory data analysis (*42, 43*). For human samples, SPEAQeasy can also perform RNA-based genotype calling with BCFtools (*44*) which can be coupled with DNA-based genotype data to identify and resolve sample swaps (**Figure 4**: downstream). Additionally, for experiments involving ERCC spike-ins (*45*), SPEAQeasy generates plots by sample to quickly visualize expected versus measured concentrations for each of 92 ERCC transcripts **Figure S1**. Thus SPEAQeasy simplifies any RNA-seq based projects from human, mouse and rat-derived data and provides a bridge to the Bioconductor universe. Furthermore, the Nextflow-based implementation allows for more experienced developers to quickly add additional steps and tools, creating a flexible and scalable RNA-seq processing pipeline.

**Figure 1.**
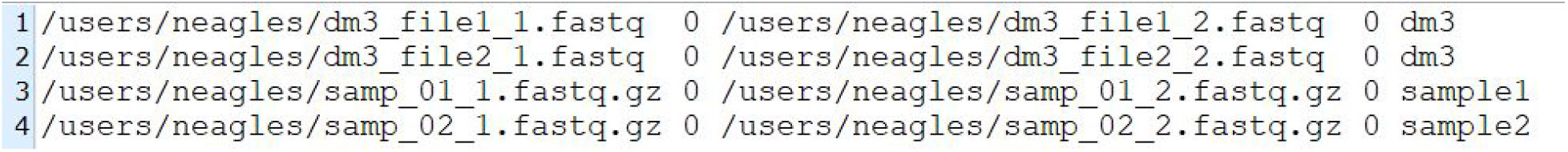
An example samples.manifest. The *samples.manifest* file for paired-end samples is composed of five tab separated columns: (1) path to the first FASTQ file in the pair, (2) optional md5 signature for the first FASTQ file in the pair, (3) path to the second FASTQ file in the pair, (4) optional md5 signature for the second FASTQ file in the pair, (5) sample ID. The first two entries use the same sample ID, which is useful when a biological sample was sequenced in multiple lanes and thus generated multiple FASTQ files. The first two pairs of FASTQ files will be merged.

**Figure 2.**
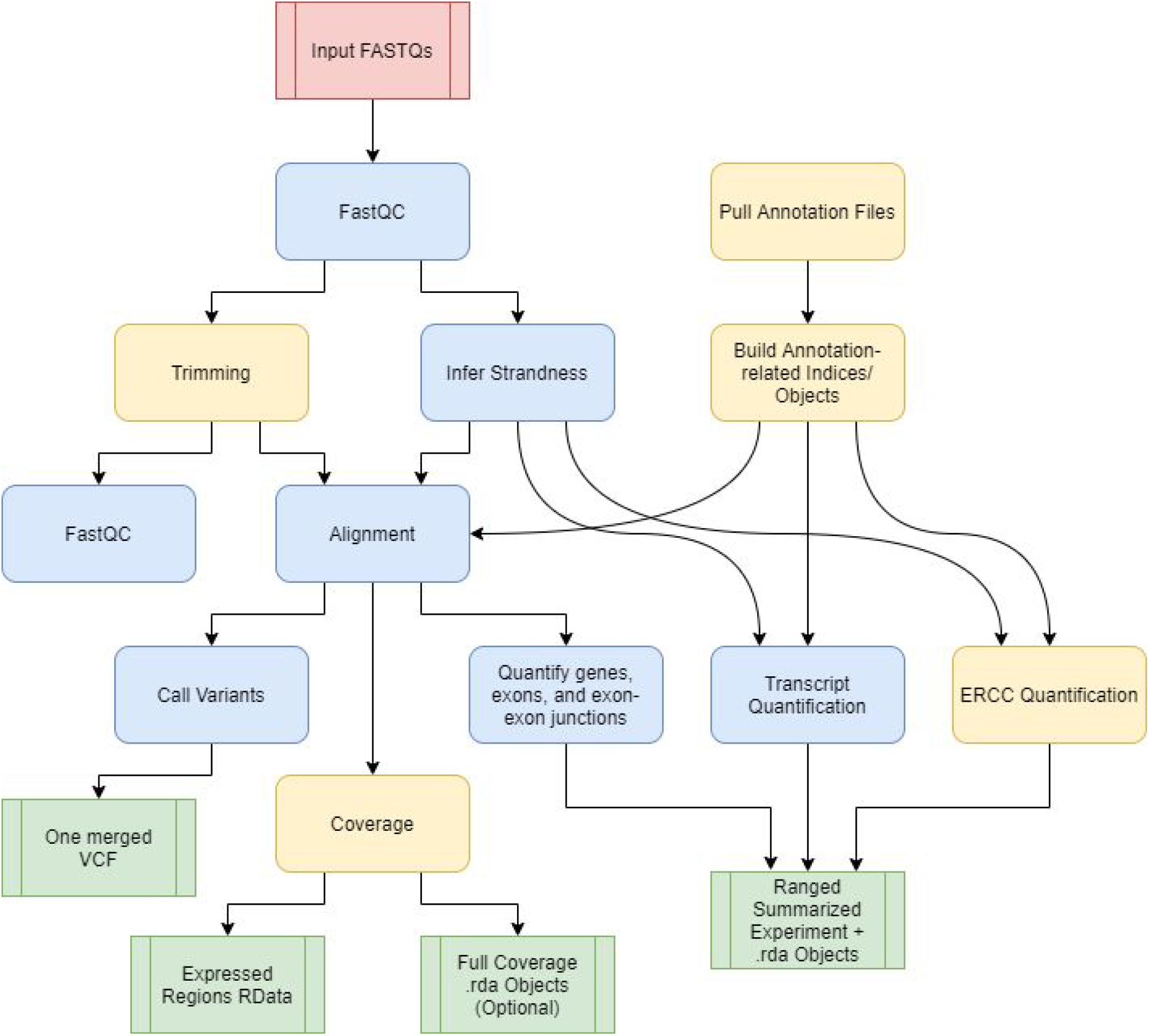
SPEAQeasy workflow diagram. A simplified workflow diagram for each pipeline execution. The red box indicates the FASTQ files are inputs to the pipeline; green coloring denotes major output files from the pipeline; the remaining boxes represent computational steps. Yellow-colored steps are optional or not always performed; for example, preparing a particular set of annotation files occurs once and uses a cache for further runs. Finally, blue-colored steps are ordinary processes which occur on every pipeline execution. The workflow proceeds downward, and each row in the diagram implicitly represents the ability for several computation steps to execute in parallel.

### Configuring SPEAQeasy

SPEAQeasy, through Nextflow (*27*), can be deployed in a variety of high-throughput computational environments such as: local machines, Sun/Son Grid Engine (SGE) compute clusters, servers that enable Docker (*46*), and cloud computing environments (*47*) such as Amazon AWS. Nextflow provides the ability to run the same code using configuration files that are specific to the computing environment at hand. To facilitate using SPEAQeasy we provide Docker containers for both the software and annotation files and a SPEAQeasy configuration file for such environments. For SGE or other clusters, SPEAQeasy can also use lmod (*48*) software modules such as the one we provide for the JHPCE SGE cluster https://jhpce.jhu.edu/. In order to use SPEAQeasy in a particular computing environment, identify the example configuration file (Methods: configuration; **Table S1**) that most resembles the setup, make a copy and edit accordingly. Our JHPCE lmod files and docker setup files provide installation instructions for researchers who wish to manually set up the software dependencies (http://research.libd.org/SPEAQeasy).

To test SPEAQeasy on a particular computer setup, first identify the “main” script appropriate for the environment. Scripts exist for execution at JHPCE or within SLURM, SGE, or local environments. A user of a SLURM-managed cluster would launch a test run of SPEAQeasy with:

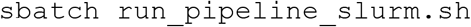

SPEAQeasy provides test samples for each combination of reference organism and strandness, which are used by default if the user does not remove the `--small_test` option and specify a directory containing the samples.manifest file with the `--input` option (Methods: test samples). While a typical test run may complete in about 15 minutes, the first execution will take significantly longer, as reference and annotation-related files must be downloaded and built for a given organism and annotation version. After successful completion, the log file `sPEAQeasy_output`. log will indicate this success at the bottom, along with details such as total run time. One can examine and become familiar with the output files from SPEAQeasy (Results: outputs), which by default are placed inside the original repository in a subfolder named `results`. Our documentation provides further detail (http://research.libd.org/SPEAQeasy/).

### Common SPEAQeasy options

Once SPEAQeasy is installed, a researcher must create a manifest file with the information about the RNA-seq samples to be processed (Methods: inputs). Next, select the “main” script written to work with the job scheduler available, if any (Methods: use cases). Within this script, a researcher may modify command options for the particular analysis. Specifically, a choice of appropriate reference genome is required to be specified with the option `--reference`, which may take values “hg19”, “hg38”, “mm10”, or “rn6”. Specify whether reads are single or paired-end with the option `--sample`, which takes values “single” or “paired”. Finally, the researcher would indicate the strandness pattern they expect all samples to obey with the option `--strand`, which may be “forward”, “reverse”, or “unstranded”. SPEAQeasy infers the actual strandness pattern present in each sample as a quality control measure (Methods: configuration; **Figure 3**: main options). See the documentation at http://research.libd.org/SPEAQeasy for further detailed options (Methods: configuration).

**Figure 3.**
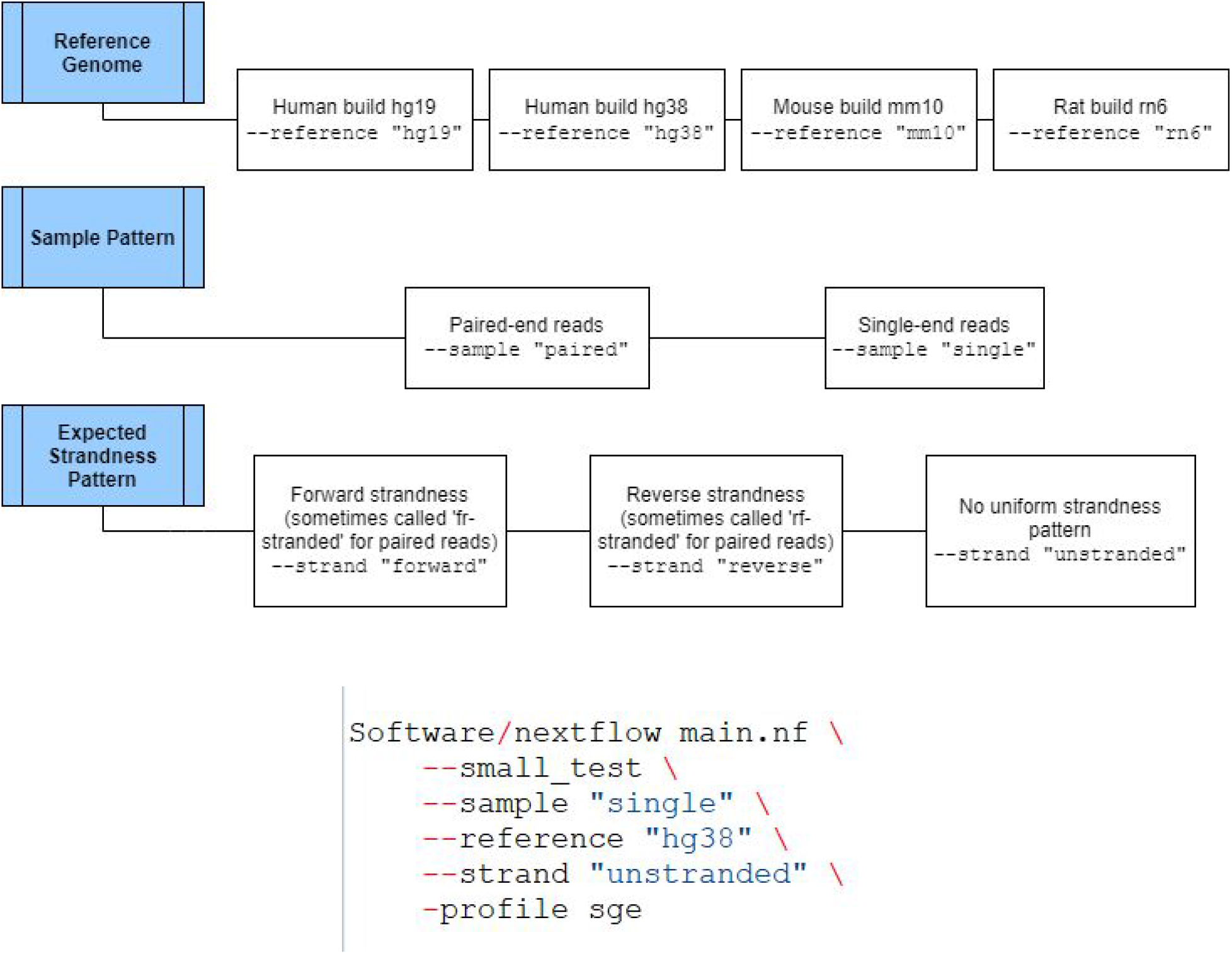
Mandatory options in the main script. The three required pieces of information the user provides are the reference genome, sample pattern, and expected strandness pattern present in all samples. The valid options are depicted horizontally to the right in this figure. On the bottom, an example of a full command is shown-in this case, a test run on an SGE scheduler without docker is also specified.

### SPEAQeasy output files

Each execution of SPEAQeasy generates a number of output files (Methods: outputs). One of the primary outputs of interest are `RangedSummarizedExperiment` R objects (*28*), which contain information about the sequence ranges, counts, and additional annotation for each feature. SPEAQeasy produces separate files for each feature type, including genes, exons, and exon-exon junctions. Because the data is packaged into `RangedSummarizedExperiment` objects, a number of Bioconductor packages can immediately be utilized to perform further analysis appropriate for a number of common use cases, starting with interactively exploring the data using tools like iSEE (*42*). A collection of quality metrics is also gathered for each sample, and saved in both an R data frame, and a comma-separated values file (**Table S2**). Users can thus assess metrics of interest at-a-glance, or utilize the information to control for covariates of interest in further analysis. Metrics include fractions of concordant, mapped, and unmapped reads during alignment, fraction of reads assigned to genes, and similar quantities.

SPEAQeasy also optionally generates bigWig coverage files for each sample, and one mean coverage file for each strand (*36*). To enable comparison between samples, coverage is normalized to 40 million mapped reads of 100 base pairs. While bigWig files may be used directly, SPEAQeasy performs an additional step to quantify coverage at genomic regions of interest. RData files are produced to describe the expressed regions (*41*), which provides a foundation for analyses involving finding differentially expressed regions.

For human samples, variant calling is performed to ultimately produce a single file for the experiment in Variant Call Format (VCF) (*49*). This file contains genotype information at a list of 740 single nucleotide variant (SNV) missense coding sites with MAF >30% (**Supplementary File 1**. Each individual typically has a unique genotype profile after variant calling, and this can be leveraged to identify mislabelled samples in conjunction with a table of identity information generated prior to sequencing, typically using a subset of the high-coverage variants in the RNA-seq data (Results: example use case involving sample swaps).

### Example use case involving sample swaps

We provide a vignette to demonstrate how SPEAQeasy outputs can be utilized to resolve sample identity issues and perform differential expression analysis (http://research.libd.org/SPEAQeasy-example/) using data from the BipSeq PsychENCODE project (*50*) which includes bulk RNA-seq data from bipolar disorder affected individuals and neurotypical controls from the amygdala and the subgenual anterior cingulate cortex (sACC). For reproducibility, the vignette walks through how to download the example data and run SPEAQeasy before performing the follow-up analysis.

First, we show how a self-correlation matrix can be constructed from user-provided genotype calls made before sequencing. The particular calls at each SNP are represented as numeric values so that an overall correlation can be computed between any two samples. The same matrix is generated from genotype calls made by SPEAQeasy (**Figure 5 A**). User-provided metadata can then be leveraged to determine if samples correlate to those of the same labelled donor, and ultimately to resolve conclusive sample swaps or drop samples with more complex identity problems. Finally, the `RangedSummarizedExperiment` objects from SPEAQeasy can be updated with these findings and metadata.

Next, explore the sources of variability in gene expression visually. First, principal component analysis is performed to assess the impact of variables such as total number of reads mapped and concordant map rate on expression (**Figure 5 B**). We also plot the first ten principal components for each individual, splitting by sex and then brain region to understand the influence of these variables on expression.

Afterward, we perform a differential expression analysis (**Table S5 A, Figure 5 C**). This involves normalizing counts with `edgeR` (*8*), forming a design matrix of interest, and controlling for heteroscedasticity in counts with `voom` (*51*). `limma` (*30*) is used to construct a linear model of expression, from which an empirical bayesian calculation can determine genes which are significantly differentially expressed. We then select genes above a particular significance threshold, in this case p<0.2, and plot expression against variables of interest. We show how to construct an expression heatmap with `pheatmap` (*52*) for top genes, with clusters labelled with covariates of interest-in this case, sex, brain region, and diagnosis status (**Figure 5 D**).

At the end, we perform a gene ontology analysis using the package clusterprofiler (*43*). The goal is to associate significantly differentially expressed genes with known functionality and biological processes. We show how to form example queries with the `compareCluster` function, and write the results to a CSV format (**Table S5 B**).

## Discussion

A number of “end-to-end” pipelines for RNA-seq are already publicly available (*19–22*). However, the majority are difficult to install or configure, require manual handling of annotation-related files, or generally lack the degree of features we have developed in SPEAQeasy (**Table S3**).

A common pipeline installation pattern involves the use of conda (*53*), where users activate and load environments where the software dependencies are installed. If conda is already available on the system, the installation process itself is typically straightforward and well-documented. However, sharing pipeline access among multiple users (e.g. in a research group/laboratory) is often nontrivial for inexperienced users, may require every individual to separately install, and this common use-case is not always documented. In contrast, SPEAQeasy provides more than one installation option, and multiple users can share a single installation instance with a single copy command: copying the main script and optionally the configuration file, which can then be modified for the individual use-case. The preferred installation method relies on Nextflow (*27*) to automatically pull pre-specified docker images at run-time unless they were previously downloaded; this approach is used in some currently-available pipelines (*19*). One of the goals of SPEAQeasy was to provide a straightforward installation method that required neither knowledge of a software/environment management tool (e.g. conda, docker, singularity, etc) nor root access permissions. Consequently, we also provide an alternative method for Linux users performed via a single command (Methods: software management):

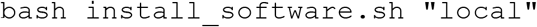

Another major focus in SPEAQeasy involves minimizing users needing to configure the pipeline for the execution environment. While many existing pipelines - in theory - support execution on a number of resource managers/ job scheduling platforms, few are pre-configured to truly leverage individual setups. For example, snakemake-based (*54*) pipelines (*21, 22*) allow specification of the total number of CPU cores to allocate, behaving identically on a local machine as on an arbitrary computing cluster. However, in practice, cluster users often must consider several other hardware resources, such as memory or disk space usage. Most notably, users of SLURM-based clusters may be charged based on specified run times of individual jobs. In the case of Nextflow and snakemake-based workflows, individual jobs are internally submitted for each pipeline component, and typically it is implicitly left to the user to worry about time specification for every component. To address this common use-case, we have written and tested configuration files for a number of environments (local execution, SGE-based clusters, SLURM-based clusters), establishing sensible defaults for variables like job run-time, memory, and disk usage.

SPEAQeasy provides other miscellaneous features we have not seen frequently or at all in other available pipelines (**Table S3**). The first involves being able to automatically handle input FASTQ samples split across more than one file. Each line in the `samples.manifest` file (Methods: inputs) specifies the path for one read or pair of reads for a sample, followed by an associated ID; for samples split across input FASTQ files just repeat the same ID for each set (line) of input files. Another feature is custom per-sample logging, which traces the exact series of commands executed, along with some additional context helpful for debugging such as relevant working directories, exit statuses, and other logging information for each process (**Figure S2**). We were motivated to implement this feature after observing how as the pipeline grew in complexity, it became increasingly necessary to understand Nextflow’s implementation details to debug execution errors. Because even a correctly written software pipeline can encounter errors when the input for a processing step is unexpectedly different or the software has a bug, we believe pipelines without specialized debugging tools become inaccessible to most users upon errors.

Software pipelines are sometimes not actively maintained. Given our interest in using SPEAQeasy ourselves (*24–26, 55*), we are actively maintaining SPEAQeasy by adapting it as new software is released for different processing steps or bugs are resolved in newer versions of the SPEAQeasy dependencies. SPEAQeasy includes an example dataset which we internally use for testing the execution as we make updates to SPEAQeasy. Given the open-source nature of SPEAQeasy and Nextflow, the SPEAQeasy code can be adapted if users are interested in switching processing tools or want to expand support to other genome references beyond `mm10, rn6, hg19` and `hg38`. The SPEAQeasy code is available on GitHub https://github.com/LieberInstitute/SPEAQeasy and https://github.com/LieberInstitute/SPEAQeasy-example, and can be expanded through interactions with users.

We anticipate that SPEAQeasy will be useful for exploring gene expression at a finer resolution, such as using exon and exon-exon junction data. The latter is powerful for exploring the un-annotated transcriptome along with base-pair coverage data (*35*). SPEAQeasy will benefit from the development of statistical and bioinformatics methods that integrate results across multiple levels of expression.

## Methods

### Overview

Pipeline execution begins with a preliminary gauge of read quality and other quality metrics, via FastQC 0.11.8 (*15*). Reads are then optionally trimmed using Trimmomatic 0.39 (*56*), and a post-trimming quality assessment is performed again with FastQC. Alignment to a reference genome is performed with HISAT2 2.1.0 (*37*), along with pseudoalignment to the transcriptome with kallisto 0.46.1 (*38*) or Salmon 1.2.1 (*39*). A combination of regtools 0.5.1 (*40*) and FeatureCounts (Subread 2.0.0) (*14*) is used to quantify genes, exons, and exon-exon junctions. At the same time, expressed regions (ERs) are optionally computed with the Bioconductor (*29*) R package derfinder (*41*). The result is a `RangedSummarizedExperiment` (*29*) object with counts information, RData files with ER information, and plots visualizing the associated data. Variant calling is also performed for human samples, using bcftools 1.10.2 (*44*) to produce a VCF file (*49*) for the experiment.

### Configuration

Usage of SPEAQeasy involves configuring two files: the “main” script and a configuration file. The “main” script contains the command which runs the pipeline, along with options specific to the input data, and fundamental choices about how the pipeline should behave. In this script, the researcher must specify if reads are paired-end or single-end, the reference species/genome (i.e. hg38, hg19, mm10, or rn6), and the expected strandness pattern to see in all samples (e.g. “reverse”). Strandness is automatically inferred using pseudoalignment rates with kallisto (*38*), and the pipeline can be configured to either halt upon any disagreement between asserted and inferred strand, or simply warn and continue. In particular, we perform pseudoalignment to the reference transcriptome using a subset of reads from each sample, trying both the *rf-stranded* and *fr-stranded* command-line options accepted by kallisto. The number of successfully aligned reads for each option is used to deduce the actual strandness for each sample. For example, an approximately equal number (40-60%) of aligned reads for each option suggests the reads lack strand-specificity and are thus “unstranded”; a large enough fold-difference between the two indicates either “reverse” or “forward “-strandness. Specifically, greater than 80% of total reads aligned must have aligned using the *rf-stranded* option to deduce a sample is “reverse”-stranded, and less than 20% to infer “forward “-strandness. We have found these cutoffs to reliably identify inaccurate `--strand` specification from the user, while not being so strict as to mistakenly disagree with correct specification. Another example command option in the “main” script controls whether to trim samples based on adapter content metrics from FastQC (*15*), trim all samples, or not perform trimming at all.

The configuration file exists for more fine-tuning of pipeline settings (such as fragment mean and standard deviation-values passed to kallisto when quantifying transcripts) and hardware resource demands for each pipeline component. Ease of use is a core focus in SPEAQeasy, and configuration files for SLURM, SGE, and local linux environments are pre-built with sensible defaults. The user is not required to modify the configuration file at all to appropriately run SPEAQeasy; however, a great degree of control and customization exists for those users who desire it.

### Inputs

A single file, called *samples.manifest*, is used to point SPEAQeasy to the input FASTQ files, and associate samples with particular IDs. It is a table saved as a tab-delimited text file, containing the path to each read (or pair of reads), optional MD5 sums, and a sample ID. Sample IDs can be repeated, which allows samples initially split across multiple files to be merged automatically (**Figure 1**). Input files must be in FASTQ format, with “.fq” and “.fastq” extensions supported, and possibly with the additional “.gz” extension for gzip-compressed files.

### Outputs

SPEAQeasy produces several output files, some of which are produced by the processing tools themselves (**Table S4**) and others by SPEAQeasy for facilitating downstream analyses (**Figure 4**). The main SPEAQeasy output files, relative to the specified `--output` directory, are:

- Under the `count_objects`/directory, `rse_gene_[experiment name].Rdata, rse_exon_[experiment name].Rdata, rse_jx_[experiment name].Rdata` and `rse_tx_[experiment name].Rdata`: these are `RangedSummarizedExperiment` objects (*29*) that contain the raw expression counts (gene & exon: featureCounts; exon-exon junctions: from regtools; transcript: either kallisto or Salmon counts), the quality metrics as the sample phenotype data (**Table S2**), and the expression feature information that depends on the reference genome used.
- Under the `merged_variants/` directory for human samples, `mergedVariants.vcf.gz`: this is a variant common format (VCF) file (*49*) with the information for 740 common variants that can be used to identify sample swaps. For example, if two or more brain regions were sequenced from a given donor, the inferred genotypes at these variants can be used to verify that samples are correctly grouped. If external DNA genotype information exists from a DNA genotyping chip, one can then verify that the RNA sample indeed matches the expected donor, to ensure that downstream expression quantitative trait locus (eQTL) analyses will use the correct RNA and DNA paired data.
- Under the `coverage/bigWigs/` directory when SPEAQeasy is run with the `--coverage` option, `[sample_name].bw` for unstranded samples or `[sample_name].forward.bw` and `[sample name].reverse.bw` for stranded samples: these are base-pair coverage bigWig files standardized to 40 million 100 base-pair reads per bigWig file. They can be used for identification of expressed regions in an annotation-agnostic way (*41*), for quantification of regions associated with degradation such as in the qSVA algorithm (*57*), visualization on a genome browser (*58*), among other uses.

**Figure 4.**
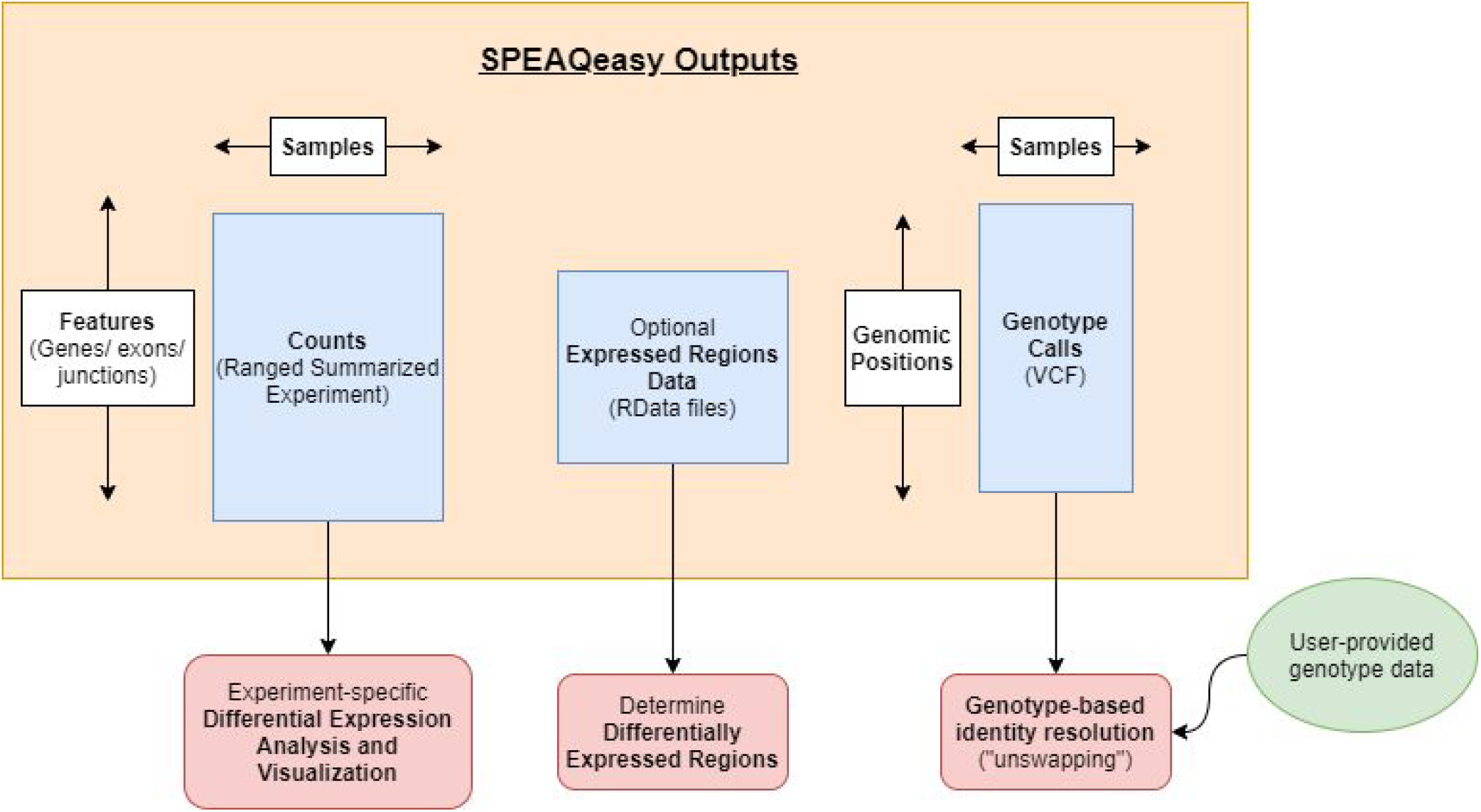
Main output files from SPEAQeasy. SPEAQeasy produces the files described in the blue boxes, as the final products of interest. Counts of genes, exons, and exon-exon junctions are aggregated into three respective R objects of the familiar `RangedSummarizedExperiment` class. This allows users to immediately follow up with a number of Bioconductor tools to perform any desired differential expression analyses. If the `--coverage` option is provided, RData files are produced to provide expression information over regions in the genome. This allows users to compute differentially expressed regions using any of a number of Bioconductor packages as appropriate for the experiment. Finally, for experiments on human samples, variants are called to ultimately produce a single VCF file of genotype calls at 740 particular SNVs. Together with genotype data recorded before sequencing the samples, one can resolve mislabellings and other identity issues which inevitably occur during the sequencing process (http://research.libd.org/SPEAQeasy-example).

**Figure 5.**
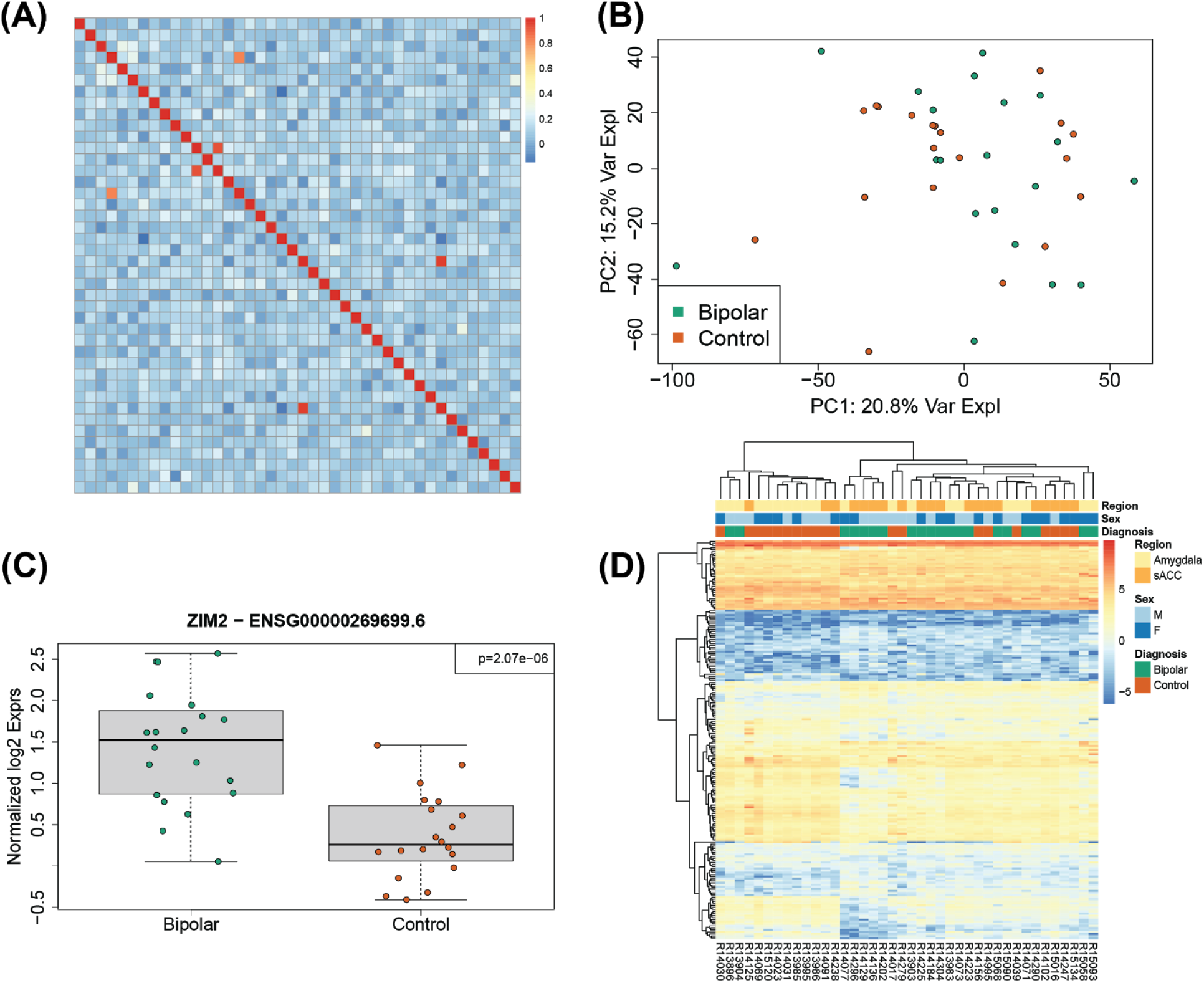
Example analysis results from applying SPEAQeasy to a subset of the BipSeq PsychENCODE dataset. **(A)** Heatmap of the spearman correlation across samples using variant information derived from the RNA-seq data produced by SPEAQeasy. Off-diagonal high correlation values indicate potential sample swaps. **(B)** Top two principal components (PCs) derived from the gene expression counts produced by SPEAQeasy colored by diagnosis. **(C)** Boxplots of the normalized log2 expression for the top differentially expressed between controls and bipolar disorder affected individuals using a subset of the BipSeq PsychENCODE data processed using SPEAQeasy. **(D)** Heatmap of the top differentially expressed genes with annotations for the brain region (amygdala or sACC), sex (male or female) and diagnosis (bipolar or control). See http://research.libd.org/SPEAQeasy-example/ for the full example analysis.

### Software Management

SPEAQeasy provides two options for managing software dependencies. If docker (*59*) is available on the system the user intends to run the pipeline, software can be managed in a truly reproducible and effortless manner. As a pipeline based on Nextflow, SPEAQeasy can isolate individual components of the workflow, called *processes*, inside docker containers. Containers describe the entire environment and set of software versions required to run a pipeline command (such as *hisat2-align*), eliminating common problems that may occur when a set of software tools (such as SPEAQeasy) is installed on a different system than it was developed. Each docker image is pulled automatically at runtime if not already downloaded (on the first pipeline run), and otherwise the locally downloaded image is used.

Because docker is not always available, or permissions are not trivial to configure, software dependencies may alternatively be locally installed. From within the repository directory, the user would run the command:

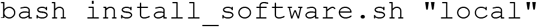

This installs each software utility from source, where available, and as a pre-compiled binary otherwise. Because installation is performed within a subdirectory of the repository, the user need not have root access for the majority of tools. However, we require that Java and Python3 be globally installed. The motivation for this requirement is that we expect most users to have these tools already installed globally, and local installation of these tools is generally advised against because of potential conflicts with other installations on the system.

Though docker and local software installation are the officially supported and recommended methods for managing software, other alternatives exist for interested users. SPEAQeasy includes a file called `conf/command_paths_long`. config, containing the long paths for each software utility to be called during pipeline execution. Users can substitute in the paths to already-installed software versions for any utility, in this file. Those familiar with Lmod environment modules (*48*) can also trivially specify in their configuration file module names to use for a particular SPEAQeasy process. However, this tends to only be a viable option for those with a diverse set of bioinformatics modules already installed.

### Annotation

SPEAQeasy is intended to be greatly flexible with annotation and reference files. By default, annotation files (the reference genome, reference transcriptome, and transcript annotation) are pulled from GENCODE (*60*) for human and mouse samples, or Ensembl (*61*) for rat samples. The choice of species is controlled by the command flag `“--reference”` in the “main” script, which can hold values “hg38”, “hg19”, “mm10”, or “rn6”. In the configuration file, simple variables control the GENCODE release or ensembl version to be used. When the pipeline run is executed, SPEAQeasy checks if the specified annotation files have already been downloaded. If so, the download is not performed again for the current or future runs. This reflects a general feature of SPEAQeasy, provided by its Nextflow base-processes are never “repeated” if their outputs already exist. The outputs are simply cached and the associated processes are skipped.

SPEAQeasy also offers easy control over the particular sequences included in the analysis-a feature we have not seen in other publically-available RNA-seq pipelines utilizing databases such as GENCODE or Ensembl. In particular, researchers are sometimes only interested in alignments/results associated with the canonical reference chromosomes (e.g. chr1-chr22, chrX, chrY, and chrMT for homo sapiens). Alternatively, sometimes extra contigs (sequences commonly beginning with “GL” or “KI”) are a desired part of the analysis as well. RNA-seq workflows commonly overlook subtle disagreement between the sequences aligned against, and sequences included in downstream analysis. SPEAQeasy provides a single configuration variable, called `anno_build`, to avoid this issue, and capture the majority of use cases. Setting the variable to “main’’ uses only the canonical reference sequences for the entire pipeline; a value of “primary” includes additional contigs’ seen in GENCODE (*60*) annotation files having the “primary” designation in their names (e.g. `GRCh38.primary_assembly.genome.fa`).

Users are not limited to using GENCODE/Ensembl annotation, however. Instead, one can optionally point to a directory containing the required annotation files with the main command option `“--annotation [directory path]”`. To specify this directory contains custom annotation files, rather than the location to place GENCODE/Ensembl files, one uses the option `“--custom_anno [label]”`. The label associates internally-produced files with a name for the particular annotation used. The required annotation files include a genome assembly fasta, a reference transcriptome fasta, and a transcriptome annotation GTF. For human samples, a list of sites in VCF format (*49*) at which to call variants is also required. Finally, if ERCC quantification is to be performed, an ERCC index for kallisto must be provided (*45*).

### Use cases

We expect that the majority of users will have access to cloud computing resources or a local computing cluster, managing computational resources across a potentially large set of members with a scheduler such as Simple Linux Utility for Resource Management (SLURM) or Sun Grid Engine / Son of Grid Engine (SGE). However, SPEAQeasy can also be run locally on a linux-based machine. For each of these situations, a “main” script and associated configuration file are pre-configured for out-of-the-box compatibility. For example, a SLURM user would open `run_pipeline_slurm.sh` to set options for his/her experiment, and optionally adjust settings in `conf/slurm.config` (or `conf/docker_slurm.config` for docker users).

In the configuration file, simple variables such as “memory” and “cpus” transparently control hardware resource specification for each process (such as main memory and number of CPU cores to use). These syntaxes come from Nextflow, which manages how to translate these simple user-defined options into a syntax recognized by the cluster (if applicable). However, Nextflow also makes it simple to explicitly specify cluster-specific options. Suppose, for example, that a particular user intends to use SPEAQeasy on an SGE-based computing cluster, but knows his/her cluster limits the default maximum file size that can be written during a job. If a SPEAQeasy process happens to exceed this limit, the user can find the process name in the appropriate config file (**Table S1**), and add the line `“clusterOptions = ‘−l h_fsize=100G’”` (this is the SGE syntax for raising the mentioned file size limit to 100G per file, a likely more liberal constraint).

We also expect a common use case would involve sharing a single installation of SPEAQeasy among a number of users (e.g. a research lab). A new user wishing to run SPEAQeasy on his/her own dataset simply must copy the appropriate “main” script (e.g. `run_pipeline_slurm.sh`) to a desired directory, and modify it for the experiment. All users then benefit from automatic access to any annotation files which have been pulled or built by the pipeline in the past, and by default share configuration (potentially reducing work in optimizing setup specific to one’s cluster). However, user-specific annotation locations or configuration settings can be chosen by simple command-line options, if preferred.

### Test samples

The test samples were downloaded from the Sequence Read Archive (SRA) or simulated using polyester (*62*), depending on the organism, strandness, and pairing of the samples. Each was then subsetted to 100,000 reads.

- Human:
  ○ Single-end, reverse: SRS7176970 and SRS7176971 (*63*)
  ○ Single-end, forward: ERS2758385 and ERS2758384
  ○ Paired-end reverse: SRS5027402 and SRS5027403 (*64*)
  ○ Paired-end forward: samples dm3_file1 and dm3_file2 from Rail-RNA (*65*); samples sample_01 and sample_02 generated with polyester (*62*)

- Mouse
  ○ Single-end, reverse: SRS7205735 and ERS3517668
  ○ Single-end, forward: all files generated with polyester (*62*)
  ○ Paired-end, reverse: SRS7160912 and SRS7160911
  ○ Paired-end, forward: all files generated with polyester (*62*)

- Rat
  ○ Single-end, reverse: SRS6431375
  ○ Single-end, forward: all files generated with polyester (*62*)
  ○ Paired-end, reverse: SRS6590988 and SRS6590989 (*66*)
  ○ Paired-end, forward: all files generated with polyester (*62*)

## Acknowledgements

We would like to acknowledge the authors of processing software tools SPEAQeasy is based upon, especially those who answered our questions on GitHub issues, support forums and emails.

## Funding

This project was supported by the Lieber Institute for Brain Development and NIH R21MH120497-01.

## Author Contribution

- N.J.E. - Conceptualization, Methodology, Software, Writing - Original Draft, Visualization
- E.E.B. - Conceptualization, Methodology, Software
- J.L. - Methodology, Software
- B.K.B. - Methodology, Software
- J.M.S. - Formal Analysis
- L.H. - Data Curation
- B.N.P. - Software
- V.L.S. - Software
- E.G-M. - Software
- I.A-O. - Methodology, Software, Project administration
- A.E.J. - Conceptualization, Methodology, Software, Writing - Review & Editing, Project administration
- L.C-T. - Conceptualization, Methodology, Software, Writing - Original Draft, Writing - Review & Editing, Project administration

## Declaration of Interests

J.L., V.L.S., E.G-M., I.A-O. were employed by Winter Genomics. All other authors have no conflicts of interest to declare.

## Supplementary Figures

**Figure S1.**
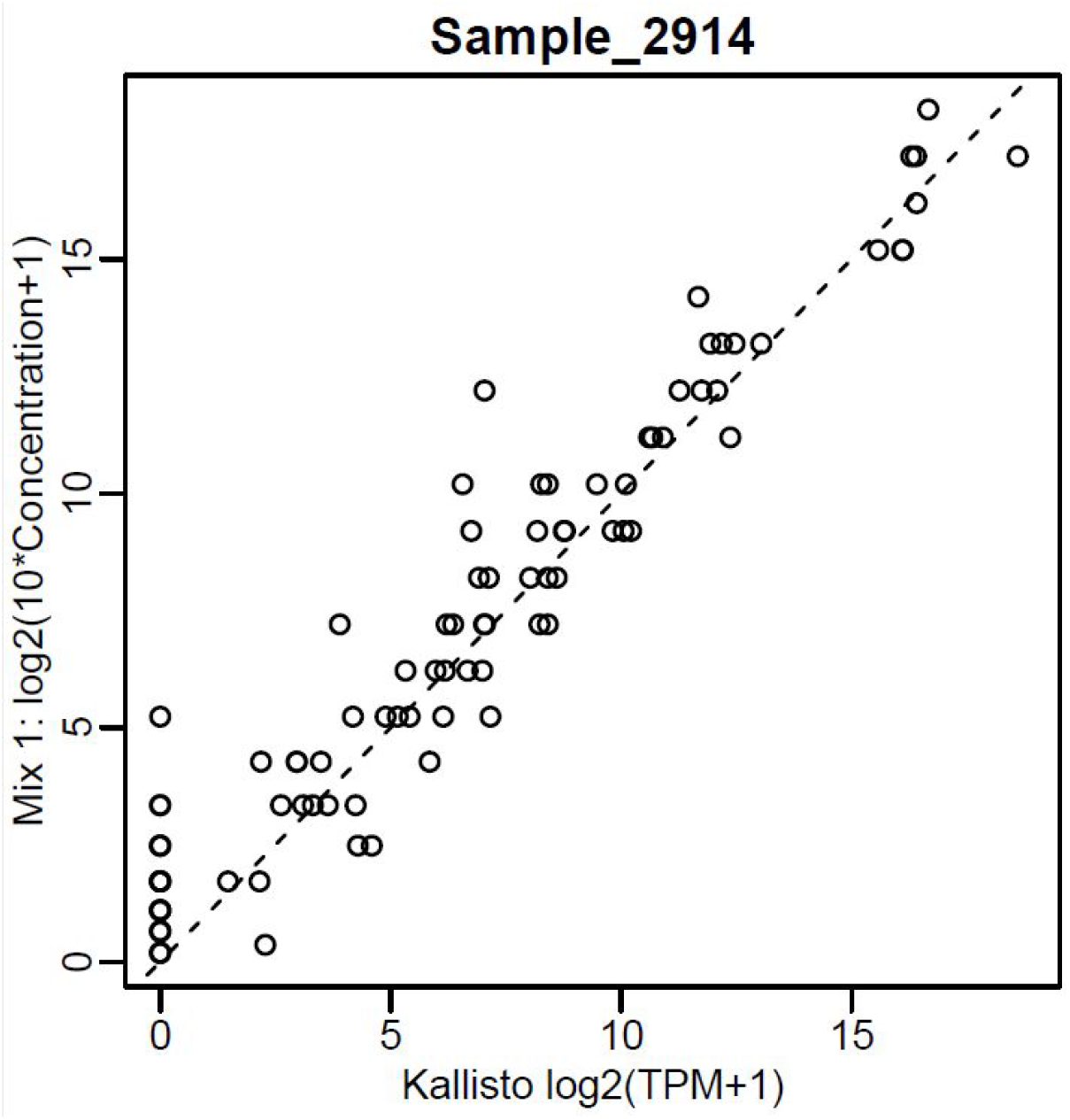
Expected vs. Actual ERCC concentration. SPEAQeasy produces plots for each sample, for easy visual comparison of expected ERCC transcript abundance with the kallisto-measured concentration.

**Figure S2.**
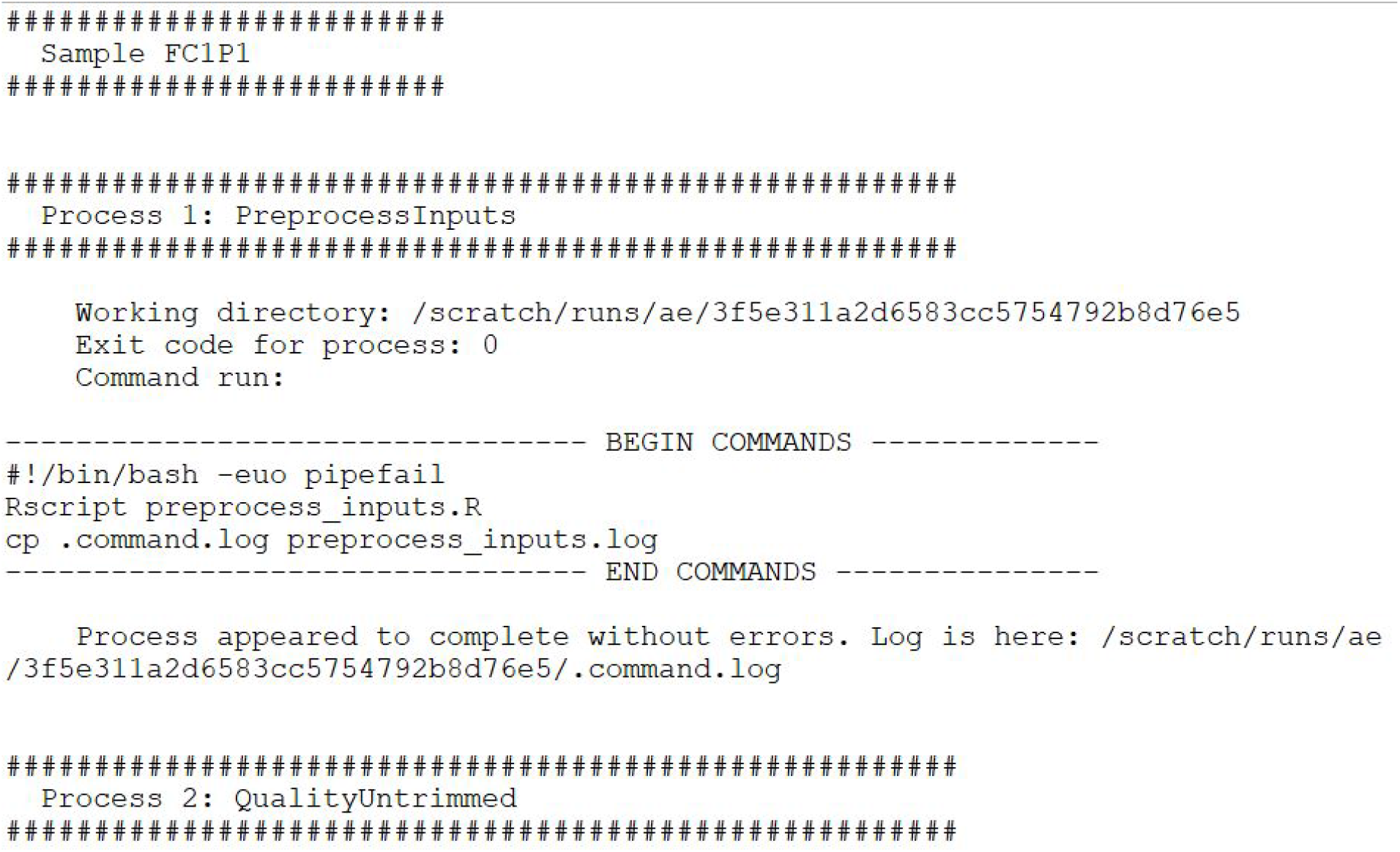
SPEAQeasy logs tracing computational steps by sample. To aid transparency and greatly simplify the source of execution errors, SPEAQeasy automatically generates logs with several pieces of information for every sample. In order of submission, the name of each Nextflow process is printed, along with (1) the working directory: where all relevant files are present, (2) the exit code: a standard indication of whether the process succeeded or how it failed, (3) a list of the specific commands run during the given process. Above is a snapshot of the top of an example log.

## Supplementary Tables

**Table S1.** Available configuration profiles. Configuration files exist under the `SPEAQeasy/conf directory`. Configuration profiles exist for SGE and SLURM clusters, as well as local execution on a Linux machine. These profiles can be customized for specific clusters, such as the JHPCE configuration file `jhpce.config`, which runs on an SGE cluster. The file a user chooses also depends on whether software dependencies are managed with docker, or are installed locally.

|  | Local installation | Installation with Docker |
| --- | --- | --- |
| Local Linux machine | local.config | docker_local.config |
| SGE scheduler | sge.config | docker_sge.config |
| SLURM scheduler | slurm.config | docker_slurm.config |
| The JHPCE cluster | jhpce.config | N/A |

**Table S2.** Quality metrics recorded in SPEAQeasy outputs. One of the major pipeline outputs is a comma-separated values (CSV) file where fields (columns) are different quality metrics, and each line (row) is associated with one sample. A list of the exact field names and their descriptions is given above.

| Metric name | Description |
| --- | --- |
| SAMPLE_ID | The name of the sample, as specified in the last column of *samples.manifest* |
| ERCCsumLogErr | If applicable, a summary statistic quantifying overall difference of expected and actual ERCC concentrations for one sample |
| trimmed | A boolean value ("TRUE" or "FALSE"), indicating whether the given sample underwent trimming |
| numReads | The number of reads present in any FASTQ files associated with the sample (after any trimming) |
| numMapped | The number of reads which successfully mapped to the reference genome during alignment |
| numUnmapped | The number of reads which did not successfully map to the reference genome during alignment |
| overallMapRate | The decimal fraction of reads which successfully mapped to the reference genome (i.e. $\text{numMapped} / \text{numReads}$) |
| concordMapRate | The decimal fraction of reads which aligned concordantly to the reference genome |
| totalMapped | The number of reads which successfully mapped to the canonical sequences in the reference genome (excluding mitochondrial chromosomes) |
| mitoMapped | The number of reads which successfully mapped to the mitochondrial chromosome |
| mitoRate | The decimal fraction of reads which mapped to the mitochondrial chromosome, of those which map at all (i.e. $\text{mitoMapped} / (\text{totalMapped} + \text{mitoMapped})$) |
| totalAssignedGene | The decimal fraction of reads assigned unambiguously to a gene, with featureCounts (14), of those in total |
| rRNA_rate | The decimal fraction of reads assigned to ribosomal RNA |

**Table S3.**
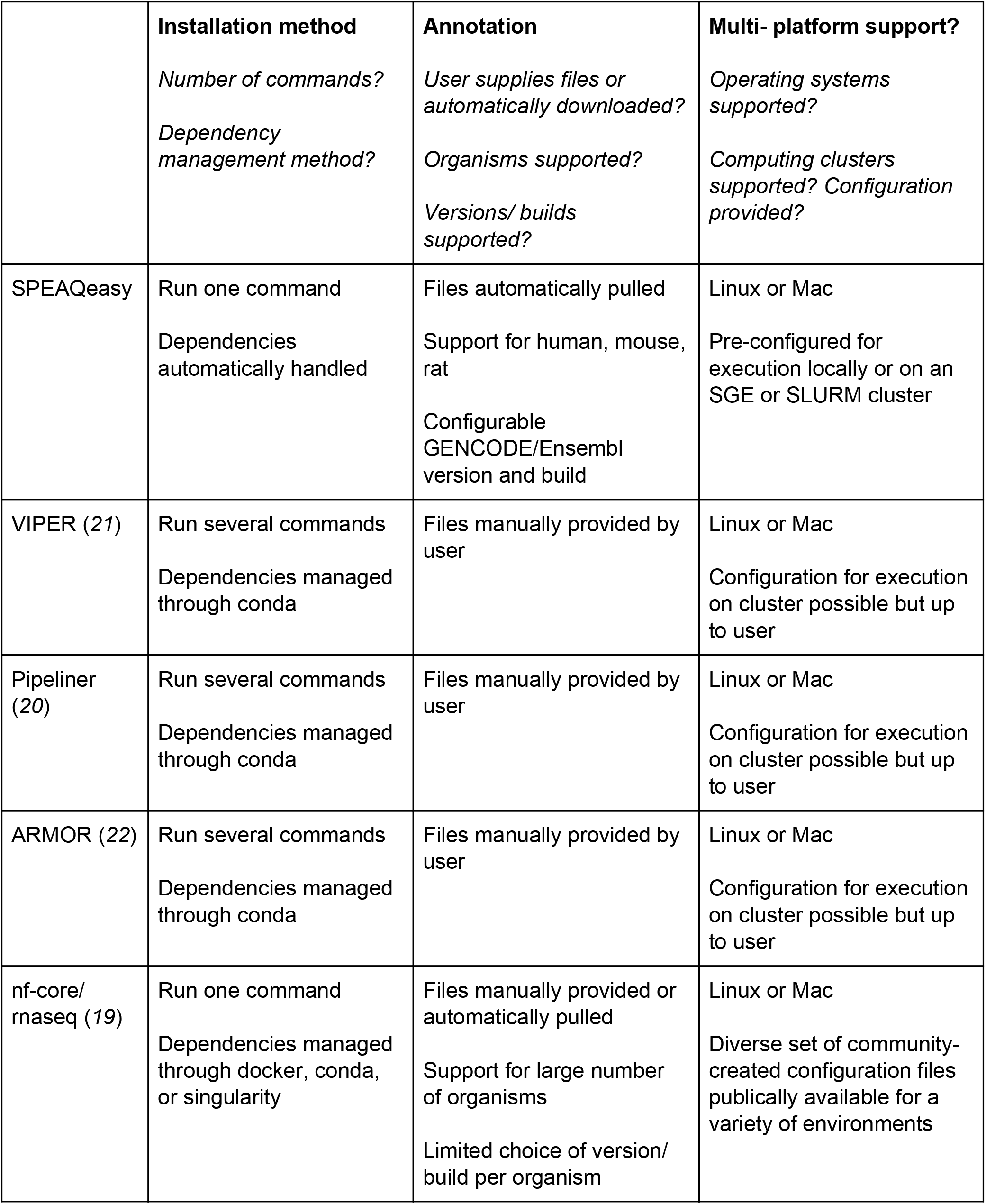
Pipeline comparison. A comparison of usage-related features among several publicly available RNA-seq pipelines.

**Table S4.** SPEAQeasy output files. Table of intermediary outputs generated by SPEAQeasy. These do not include the major output files of interest (**Figure 4**), but other miscellaneous outputs from each processing step. In the Filename column, brackets denote one or more values dependent on a relevant variable; for example, the files `[sample_name]_process_trace.log` refer to a set of several files, each named distinctly according to the sample associated with the particular file. An asterisk represents a wildcard matching more than one file, when individual file names may depend on the experiment. For example, `[sample_name]_trimmed*.fastq` could refer to `sample1_trimmed_1.fastq` and `sample1_trimmed_2.fastq`. The next columns provide the directory containing each given file, relative to the output folder, and a description of the files’ content, respectively.

| Filename | Directory | Description |
| --- | --- | --- |
| ercc_spikein_check_mix1.pdf | count_objects / | A plot comparing the Mix1 expected concentration (https://tools.thermofisher.com/content/sfs/manuals/cms_095046.txt) against the observed counts from Kallisto (38). |
| rawCounts_[experiment_name]_n[num_samples].rda | count_objects / | An R data file containing a number of matrices/ data frames containing raw gene, exon, exon-exon junction, and transcript counts. A data frame of quality metrics is also included (**Table S2**). |
| read_and_alignment_metrics_[experiment_name].csv | count_objects / | A comma-separated values file of quality metrics (**Table S2**). |
| rpkm_counts_[experiment_name]_n[num_samples].rda | count_objects / | The same data as in rawCounts_[experiment_name]_n[num_samples].rda, but normalized as reads-per-kilobase-million (RPKM). |
| [sample_name]_[annotation_version]_Exons.counts | counts/ | Exon counts for each sample individually, as reported by featureCounts (14) |
| [sample_name]_[annotation_version]_Exons.counts.summary | counts/ | A summary of key information generated from quantifying exons on a particular sample with featureCounts (14) |
| [sample_name]_[annotation_version]_Gene.counts | counts/ | Gene counts for each sample individually, as reported by featureCounts (14) |
| [sample_name]_[annotation_version]_Gene.counts.summary | counts/ | A summary of key information generated from quantifying genes on a particular sample with featureCounts (14) |
| [sample_name]_junctions_primaryOnly_regtools.bed | counts/junction/ | A file in BED format (58) describing junctions determined by regtools (40) run on primary alignments for a given sample |
| [sample_name]_junctions_primaryOnly_regtools.count | counts/junction/ | A text file describing the junction ranges and providing raw counts of hits for each junction, as output by regtools (40) |
| `[sample_name].bam` and `[sample_name].bam.bai` | `counts/junction/primary_alignments/` | The subset of primary alignments for a given sample, in BAM format (67), along with an index for each BAM. |
| `mean.forward.bw` and `mean.reverse.bw` | `coverage/mean/` | bigWig files (36) containing coverage information averaged across all samples in the experiment, and split by strand. |
| `[sample_name].[strand].wig` | `coverage/wigs/` | Wiggle files (36) containing coverage information for a given sample and strand. |
| `abundance.tsv` | `ERCC/[sample_name]/` | The output from running Kallisto (38) against the 92 ERCC RNA Spike Mix (https://www.thermofisher.com/order/catalog/product/4456740) sequences for each sample. |
| `region_cuts_[strand].Rdata` | `expressed_regions/` | An R object called `region_cuts` that is a list with one element per cutoff that contains a `GenomicRanges::GRanges()` object with the expressed regions across all chromosomes. |
| `region_cuts_raw_[strand].Rdata` | `expressed_regions/` | An R object called `region_cuts_raw` that is a list with one element per chromosome, then a nested element per cutoff used for identifying expressed regions using `derfinder::findRegions()` (41). |
| `region_info_[strand].Rdata` | `expressed_regions/` | An R object called `regInfo` that is a `data.frame` with the columns: `cutoff`, `n`, `mean`, and `sd`. This table summarizes the number, mean width, sd of the width for the expressed regions identified at each cutoff. |
| `region_info_[strand].pdf` | `expressed_regions/` | A PDF with a few exploratory plots made using `regInfo` that evaluate how the cutoff for identifying the expressed regions affects the number of ERs, their mean width, the sd of their width. These are the visualization plots recommended for choosing a cutoff as described on the `derfinder` manuscript (41) at Figure S4 (https://github.com/leekgroup/derSuppleme |
|  |  | nt/blob/gh-pages/derfinder_supplement.pdf)<br>. |
| [trim_status]/[file_name]/* | fastQC/ | Outputs from FastQC (15). Here `trim_status` indicates when FastQC was performed: `Untrimmed` is before trimming, and `Trimmed` is after. `file_name` contains the sample name, and if applicable, the mate number. |
| [sample_name]_hisat_out.sam | HISAT2_out/ | The main alignment output from HISAT2 (37) in SAM format (67) |
| [sample_name]_align_summary.txt | HISAT2_out/ | The text-based alignment summary from Hisat2 (37). Note that metrics from these files are aggregated for the experiment, and so users likely will not need to check or process the original files manually (Table S2). |
| [sample_name]_accepted_hits.sorted.bam<br>and<br>[sample_name]_accepted_hits.sorted.bam.bai | HISAT2_out/sam_to_bam/ | Alignments from Hisat2 (37) where unmapped reads are filtered out, and the resulting BAM file (67) is coordinate-sorted and indexed. |
| [sample_name]_strandness_pattern.txt | infer_strandness/ | A text file containing the inferred strandness pattern for each sample. These files are primarily for internal use by the pipeline, and it is recommended to check the output file `samples_complete.manifest` to quickly view strandness patterns for all samples. |
| samples_complete.manifest | infer_strandness/ | A version of the input `samples.manifest`, with an additional column listing the inferred strandness pattern for each sample |
| abundance.h5 | kallisto_tx/[sample_name]/ | Abundance estimates, bootstrap estimates, run metadata, and transcript length from Kallisto, saved into an HDF5 binary file. |
| [sample_name]_abundance.tsv | kallisto_tx/[sample_name]/ | Plain-text abundance estimates from Kallisto across the reference transcriptome. |
| run_info.json | kallisto_tx/[sample_name]/ | Kallisto run metadata in javascript object notation. |
| [sample_name]_process_trace.log | logs/ | A SPEAQeasy-generated log tracing the processes and associated commands run for each sample. This is intended to help users quickly determine the source of any errors during pipeline execution (**Figure S2**). |
| [sample_name]_trimmed*.fastq | trimming/ | Trimmed FASTQ files, if applicable, from Trimmomatic (56) |

**Table S5. SPEAQeasy-example differential expression and gene ontology results. (A)** Differential expression results using the subset of BipSeq data analyzed in http://research.libd.org/SPEAQeasy-example/. **(B)** Gene ontology enrichment results from the genes with a p-value < 0.005 in the differential expression results between bipolar disorder affected individuals and neurotypical controls.

**Supplementary File 1. SNVs supplementary BED files.** The common SNVs used for sample identification are stored in the BED files `common_missense_SNVs_hg19.bed` and `common_missense_SNVs_hg38.bed`.

## Notes

http://research.libd.org/SPEAQeasy/

http://research.libd.org/SPEAQeasy-example/

